# The peculiar property of pia mater on the prediction of acute subdural hematoma

**DOI:** 10.64898/2026.06.24.733734

**Authors:** Chengbin Li, Svein Kleiven, Zhou Zhou

## Abstract

Acute subdural hematoma (ASDH) is a prevalent injury with high mortality and morbidity, often resulting from bridging vein (BV) disruption secondary to cortical relative motion. As a thin membrane enveloping the brain surface and anchoring BVs, the pia mater is hypothesized to play a critical mechanical role in cortical response and hence ASDH pathogenesis. Finite element (FE) head models are valuable tools to predict ASDH occurrence during impacts. However, the pia mater is often represented as an elastic material in existing FE head models, despite experimental evidence reporting its nonlinear mechanical behavior. In this study, both linear (Young’s modulus of 11.5 MPa) and nonlinear (the stress-strain curve derived from pial tension tests) material models of the pia mater were implemented in one FE head model. The models were subjected to three experimental impact loadings, one of which was known to cause ASDH and two of which were not. Results demonstrated that, across all simulated impacts, the model with nonlinear pia mater properties predicted larger cortical displacements and BV responses than the linear model. For the impact with known ASDH occurrence, the predicted BV strain was 0.17 for the nonlinear model and 0.094 for the linear model, with only the former approaching the reported rupture strain range of the BV-superior sagittal sinus complex (0.29 ± 0.13). These findings verified the mechanical importance of the pia mater in cortical responses and hence the prediction of ASDH, suggesting that conventional linear pia modeling might over-constrain cortical motion, leading to underestimation of BV strain and ASDH risk. The current study supported the adoption of experimentally derived nonlinear pia mater properties in FE head models to improve the reliability of ASDH prediction.

## 1. Introduction

Acute subdural hematoma (ASDH) is one of the most lethal forms of head trauma (Wilberger et al. 1991), with falls and traffic collisions being the common injury scenarios (Karibe et al. 2014, Antona-Makoshi et al. 2018, Lee et al. 2018, Kapapa et al. 2024, Meng et al. 2026). The disruption of the bridging vein (BV) secondary to the relative motion between the brain cortex and skull has been considered a major cause of ASDH, in addition to brain contusion and laceration of the cerebral vessels (Famaey et al. 2015, Zhou et al. 2020). In Sweden, for instance, ASDH accounts for around 50% of all brain injuries when excluding concussions (Kleiven et al. 2003, Pedersen et al. 2015). Despite modern advancements in critical care and intensive treatment, ASDH is still associated with notably high mortality rates, which have been reported to be up to 90% (Seelig et al. 1981, Vega et al. 2017, Jost 2022, Basilio et al. 2024). Given the urgency of this crisis, efforts to advance a fundamental understanding of ASDH are crucial for developing more effective prevention strategies.

Finite element (FE) head models are valuable tools to predict ASDH risk during head impacts. As computational surrogates, FE head models can offer detailed information on intracranial responses, such as the relative cortical motion between the brain cortex and the skull. Thus, reliable brain-skull interface modeling, containing the dura, arachnoid, and pia mater, along with cerebrospinal fluid (CSF) in the subarachnoid space, is required to improve ASDH predictions (Zhou et al. 2019a, 2020). Approaches in different aspects of the brain-skull interface have been widely investigated. These approaches include modeling contact methods, such as utilizing elastic springs (Mazumder et al. 2013), tied contact (Mao et al. 2013), tie-break contact (Takhounts et al. 2003), sliding-only contact (Kleiven 2007, Yang et al. 2022), and fluid-structure interaction (Zhou et al. 2019a, b, 2020) to connect the brain cortex to the skull or the CSF, conducting parameter studies on CSF material properties (Chafi et al. 2009). However, the mechanical role of the pia mater has often been neglected (Aimedieu et al. 2004, Ozawa et al. 2004).

The cranial pia mater, as the innermost and thinnest meningeal layer, serves as the final protective barrier for the brain (Ommaya 1968, Betsholtz et al. 2024). Structurally, it consists of a thin layer of leptomeningeal cells connected by gap junctions and occasional desmosomes (Weller et al. 2018). Assumed to be an initially loose and unordered fiber network, the pia mater exhibits a nonlinear mechanical response under stretch: an initial gradual increase in force as unaligned fibers orient themselves, a subsequent steep phase corresponding to the direct stretching of these newly ordered fibers, and a final progressive upward deviation indicating the structural deterioration within the membrane (Aimedieu et al. 2004). Despite the experimental evidence of the distinctly nonlinear mechanical properties of the pia mater (Aimedieu et al. 2004), the majority of current FE head models still utilize simplified linear elastic models, with Young’s modulus ranging from 1.1 to 12.5 MPa (Kleiven 2002, Kimpara et al. 2006, Zhang et al. 2011, Mao et al. 2013, Ji et al. 2015, Wu et al. 2019, Yang et al. 2022, Bahreinizad et al. 2025). Only a handful of studies have implemented accurate nonlinear material behaviors based on experimental data (Ho et al. 2009, Zhou et al. 2019b, Atsumi et al. 2021, Li et al. 2021). However, none of these studies isolated the effect of pial constitutive model choice on ASDH predictability.

The present study aimed to investigate the biomechanical influence of pial material modeling on the prediction of ASDH. Two pia mater modeling strategies, incorporating linear and nonlinear material properties, were implemented in an FE head model. Both models were subjected to experimentally determined loading conditions that were known to cause ASDH or not. It was hypothesized that representing the pia mater with experimentally derived nonlinear material properties together with realistic brain-skull interface modeling would improve the prediction accuracy of ASDH in the FE head model. This study highlights the importance of pia mater modeling and provides guidance for improving brain-skull interface modeling and ASDH prediction in FE head models.

## 2. Methods

### 2.1 Finite element head model

The current investigation employed an FE head model originally developed by Kleiven (2007) at the KTH Royal Institute of Technology (i.e., the KTH head model) in LS-DYNA. The KTH head model included the major components of the human head, which were represented using different element types. Specifically, the scalp, skull, brain, subarachnoid CSF, ventricles, a simplified neck with the extension of the spinal cord (Figure 1A), and deep brain regions (i.e., corpus callosum, thalamus, midbrain, and brainstem in Figure 1B) were modeled as solid elements, while the dura mater, falx, and tentorium were represented using shell elements. The brain was assumed to be an isotropic and homogeneous structure and modeled as a hyper-viscoelastic material (Zhou et al. 2025). The dura mater, falx, and tentorium were modeled as simplified rubber with an experimentally derived stress-strain curve (Bylski et al. 1986, Li et al. 2017), while the material properties of the pia mater were described in section 2.2. Further details regarding the geometrical discretization, material representation, element formulation, and experimental validation of the model are available in previous studies (Kleiven 2006, Kleiven 2007, Zhou et al. 2018, Zhou et al. 2019b).

**Figure 1.**
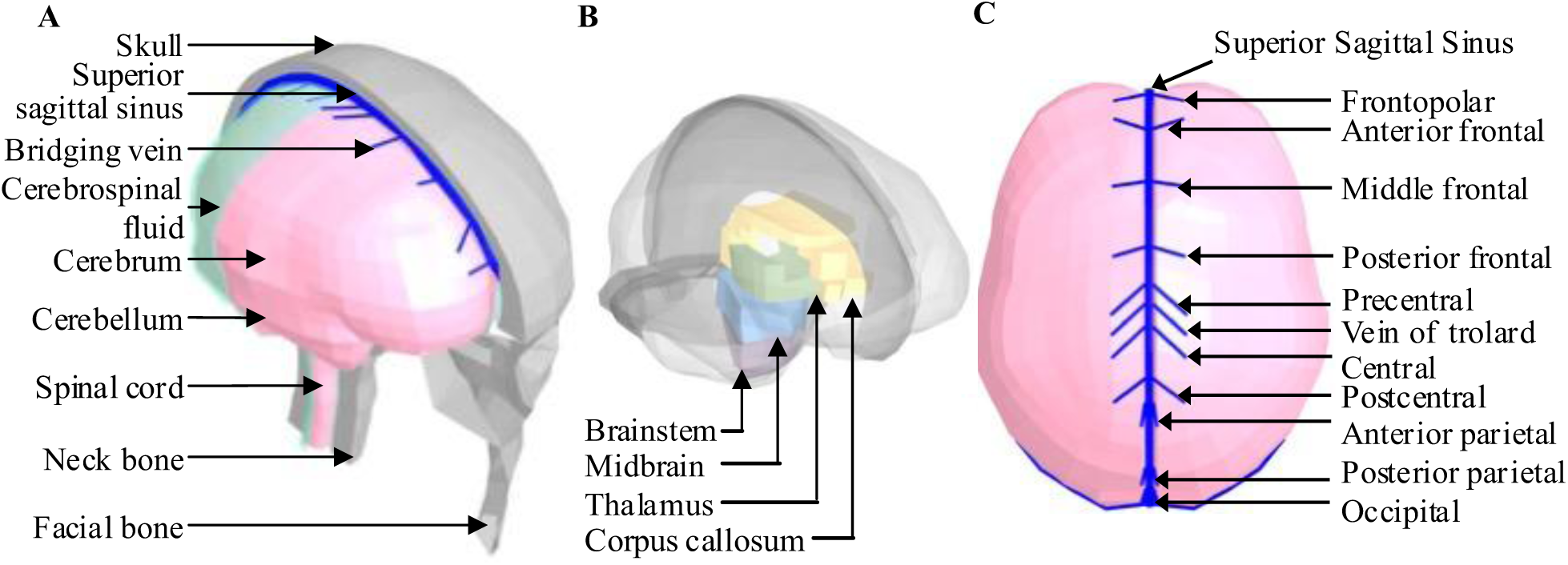
(A) Isometric view of the finite element model of human head with the skull open to expose the brain; (B) Isometric view of the deep brain structures; (C) Superior view of the brain showing the 11 pairs of the bridging veins.

Of relevance to the current study, 11 pairs of the largest parasagittal BVs were modeled as beam elements to simulate the discrete mechanical tethering between the cerebral cortex and the superior sagittal sinus (Figure 1C). The entry angles and lengths of each BV were defined according to the anatomical characterizations established by Oka et al. (1985), as detailed in Table 1. Mechanically, these elements were assigned a tension-only constitutive formulation to reflect their physiological load-bearing limitations, and a uniform tensile stiffness of 1.9 N/mm was applied to all BV elements based on the data from Lee et al. (1989).

**Table 1.** Summary of bridging veins (BVs)’ angle between superior sagittal sinus and BV elements and length of BV elements in the FE head model.

| Bridging veins | Angle (Degree) | Length (mm) |
| --- | --- | --- |
| Frontopolar | 85.9 | 12.7 |
| Anterior frontal | 124.3 | 15.6 |
| Middle frontal | 78.2 | 10.3 |
| Posterior frontal | 78.8 | 13.6 |
| Precentral | 49.5 | 18.3 |
| Vein of trolard | 47.7 | 18.2 |
| Central | 47.5 | 18.3 |
| Postcentral | 46.8 | 17.3 |
| Anterior parietal | 12.4 | 11.7 |
| Posterior parietal | 15.0 | 17.6 |
| Occipital | 5.6 | 16.8 |

### 2.2 Pia mater modeling

To assess the influence of pia mater modeling on brain cortical responses, two pial modeling strategies were implemented in the KTH head model. The first one represented the pia mater as a linear elastic material, as has been commonly used in existing head models (Kleiven 2002, Kimpara et al. 2006, Zhang et al. 2011, Mao et al. 2013, Ji et al. 2015, Wu et al. 2019, Yang et al. 2022, Bahreinizad et al. 2025). In the current study, the Young’s modulus of the linear pial material was set to 11.5 MPa, consistent with the original KTH head model (Kleiven 2007) and another counterpart (Horgan et al. 2003). In contrast, uniaxial tensile experiments on isolated cranial pia mater samples by Aimedieu et al. (2004) showed distinctly nonlinear mechanical responses. Specifically, the experimental force-displacement curve showed an initial toe region characterized by increasing stiffness, followed by a quasilinear elastic region prior to progressive deterioration and rupture. Motivated by the experimental observation, the second approach represented the pia mater using a nonlinear simplified rubber material (i.e., *MAT_SIMPLIFIED_RUBBER), similar to the approach of Ho et al. (2009) and Zhou et al. (2019b). The corresponding stress-strain curve of the nonlinear pial material was converted from the force-displacement curves measured by Aimedieu et al. (2004), assuming a pial thickness of 0.1 mm. These two models were hereafter referred to as the linear model and the nonlinear model, respectively.

In both models, the pia mater was discretized using a continuous layer of membrane shell elements that shared consistent nodes with the exterior surface of the brain (Figure 2A). A penalty-based contact was defined between the pia mater and the inner surface of the subarachnoid CSF that permitted sliding in the tangential direction and delivered tension and compression in the radial direction. The subarachnoid CSF was modeled as an elastic fluid with a bulk modulus of 2.1 GPa (Zhou et al. 2025). The localized motion responses of these two models were validated against available post-mortem human subject experiments by Alshareef et al. (2021), and a comparison of the validation results between the linear and the nonlinear model is presented in Appendix A.

**Figure 2.**
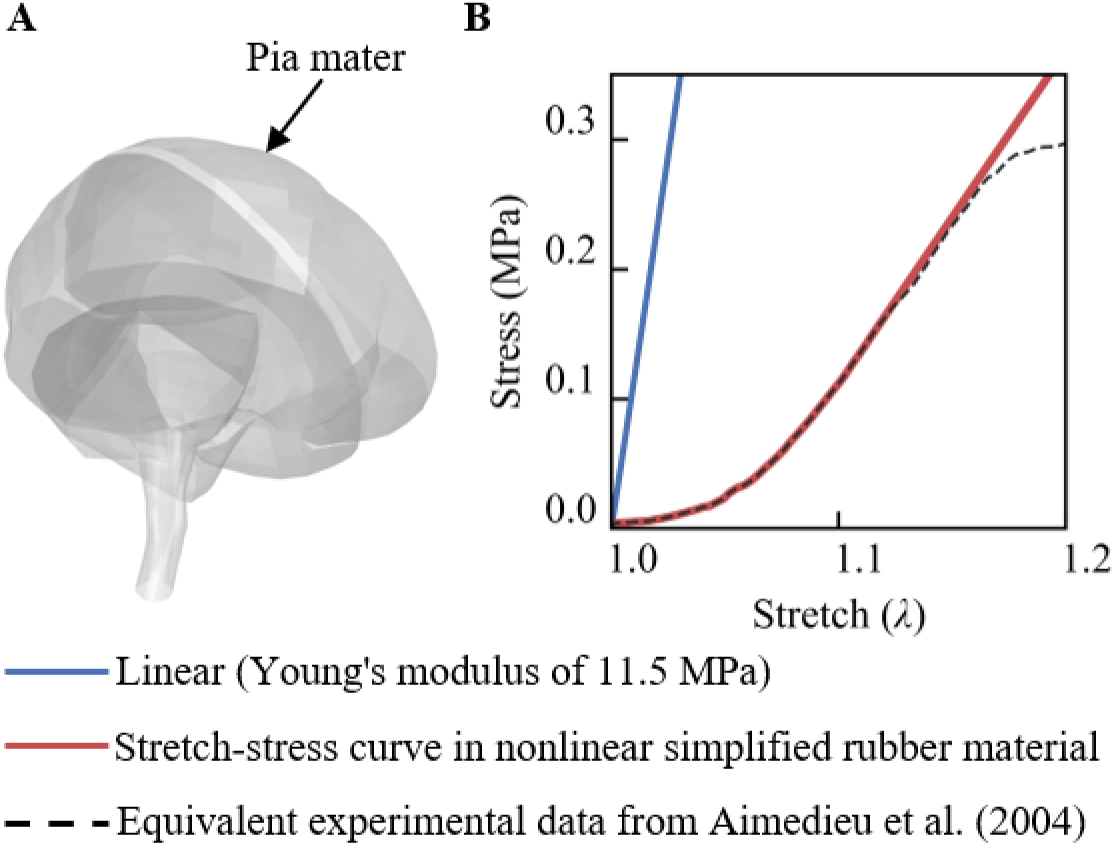
(A) Isometric view of pia mater (shown semi-transparent) modeled as shell elements with a thickness of 0.1 mm; (B) Stress–stretch curves of the linear (Young’s modulus of 11.5 MPa) and nonlinear simplified rubber material for pial modeling, compared with equivalent experimental data from Aimedieu et al. (2004).

### 2.3 Loading condition and data analysis

To analyze the influence of pia mater modeling strategies on the prediction of ASDH occurrence, whole-head impact simulations were performed. The impact loading was derived from the study by Depreitere et al. (2006), in which the unembalmed cadavers were exposed to occipital impacts with the acceleration profiles and autopsy examination (with and without BV disruption) available. In the present study, three representative impacts (Table 2) were simulated by the models with linear and nonlinear pial material. Of the three selected experiments, the loading profile of one impact resulting in BV rupture (translational acceleration peak: 450 g; rotational acceleration peak: 26.2 krad/s²) was exemplified in Figure 3.

**Figure 3.**
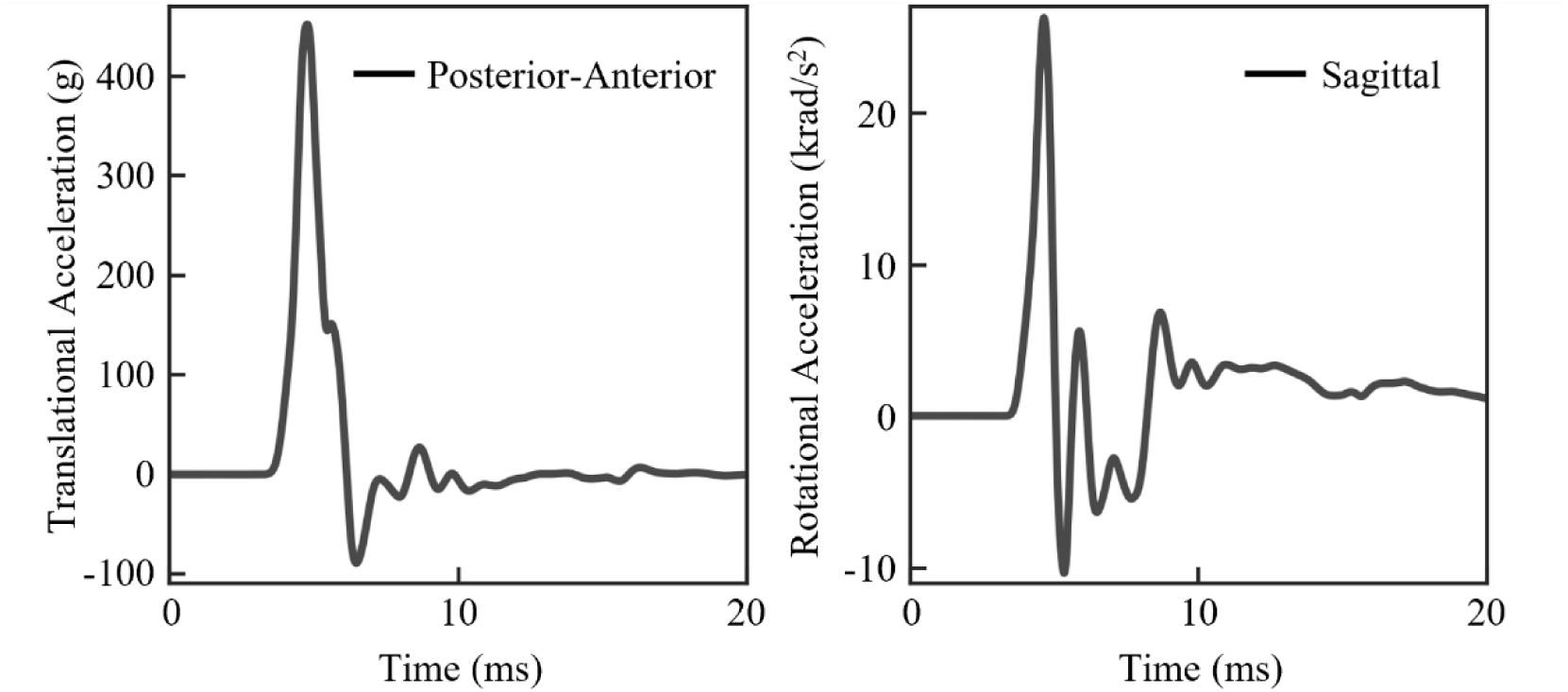
Head model loading condition for case 21-2_2 with detected bridging vein rupture.

**Table 2.** Experimental kinematics and bridging vein rupture conditions of three representative cases.

| Case | Peak rotational acceleration (rad/s<sup>2</sup>) | Peak translational acceleration (g) | Bridging vein rupture |
| --- | --- | --- | --- |
| 21-2_2 | 26262 | 450 | Yes |
| 25-2_1 | 7312 | 216 | No |
| 25-2_2 | 7975 | 202 | No |

For each simulation, the impact pulses were prescribed to a node located at the center of gravity of the FE head model, and this node was attached to the rigid skull. All simulations were solved by LS-DYNA (version 16.0 double precision). For a simulation of 30 ms, it took approximately 30 minutes to solve using 16 central processing units on Linux. The cortical motion (i.e., the nodal displacement of cerebral surface elements relative to the skull) and BV strain were used to assess the occurrence of ASDH, following the strategies in previous studies (Kleiven 2003, Zhou et al. 2019a, b). In addition, brain strain, with particular focus on the cerebral cortex, was extracted to examine the effect of the pial material on deformation responses.

## 3. Results

The material properties of the pia mater affected brain displacement responses. For the case of 21-2_2, both models predicted relatively large brain-skull relative motion (characterized by the red area in Figure 4A) at the site close to the frontal skull base. When focusing on the cortical area that is more relevant to the prediction of ASDH, only the nonlinear model with experimentally based properties predicted large cortical relative motion at the superior parietal and frontal regions. Across all three simulated impacts, the nonlinear model consistently predicted larger cortical relative motion peaks than the linear model (Figure 4B). In case 21-2_2, the nonlinear model predicted a maximum cortical relative motion of 7.72 mm, whereas the linear model predicted a much lower peak value of 1.97 mm. Similar trends were observed for the two loadings without detected BV rupture, i.e., 3.32 mm for the nonlinear model vs. 1.17 mm for the linear model in case 25-2_1 and 3.32 mm for the nonlinear model vs. 0.93 mm for the linear model in case 25-2_2.

**Figure 4.**
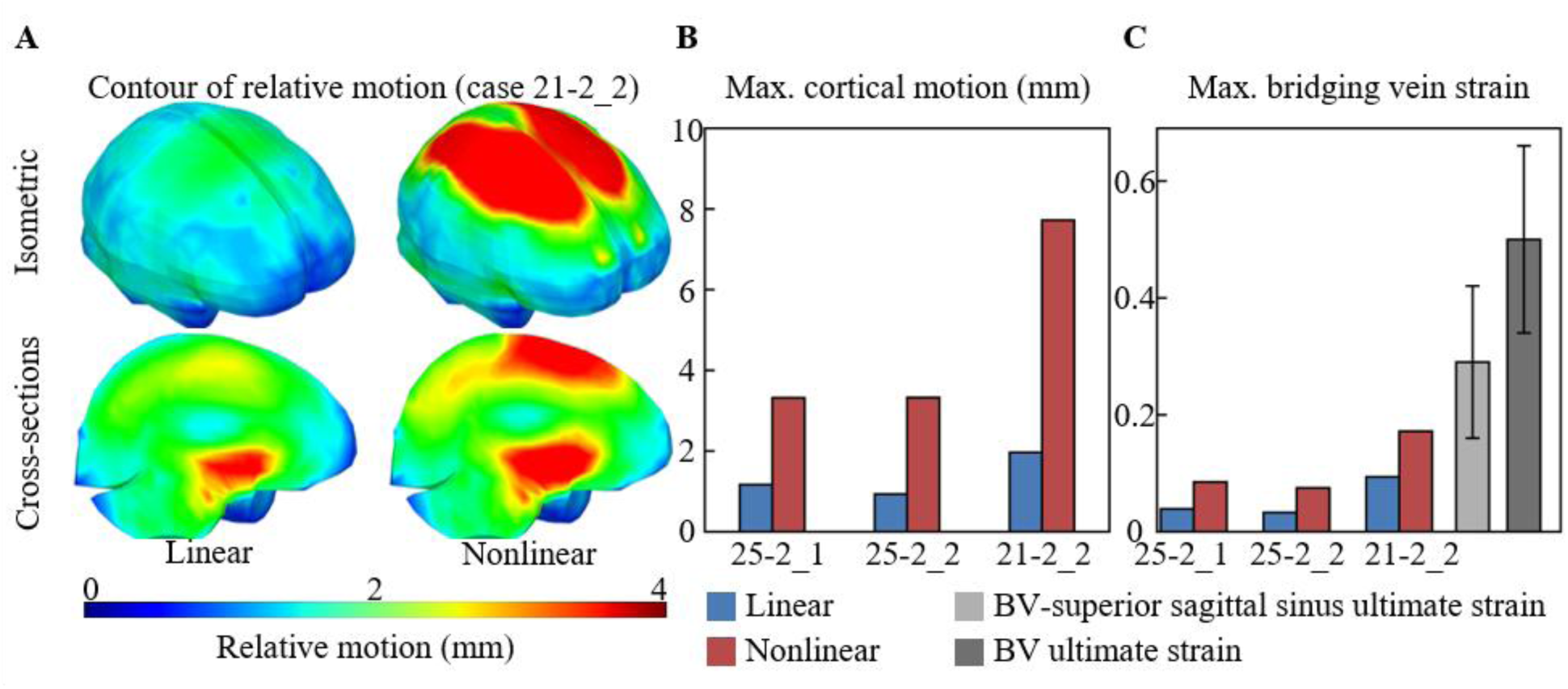
Influence of the pial modeling (linear vs. nonlinear) on the cortical relative motion and bridging vein (BV) strain in three simulated impacts (i.e., cases 25-2_1, 25-2_2, and 21-2_2). (A) Isometric view (upper) and sagittal cross-sections (lower) of cortical relative motion distribution (case 21-2_2); (B) Maximum value of the cortical relative motion; (C) Maximum value of BV strain plotted together with the ultimate strain of the BV-superior sagittal sinus complex (Monea et al. 2014), and BV ultimate strain (Lee et al. 1989).

The estimated maximum BV strains were consistently higher in the nonlinear model than in the linear counterpart (Figure 4C). Peak BV strains for the nonlinear vs. linear models were 0.17 vs. 0.094 (case 21-2_2), 0.085 vs. 0.039 (case 25-2_1), and 0.075 vs. 0.033 (case 25-2_2). When compared with the ultimate strain range (0.29 ± 0.13) of the BV-superior sagittal sinus (BV-SSS) complex (Monea et al. 2014), a possible ASDH occurrence was predicted in case 21-2_2 only when the pia mater was simulated as a nonlinear material. None of the predicted BV strains fell within the range of the BV ultimate strain (0.50 ± 0.16) (Lee et al. 1989). In case 21-2_2, the peak BV strain was located on the precentral veins in the nonlinear model and was consistent with the experiment by Depreitere et al. (2006), which reported that bridging vein ruptures invariably occurred in the rolandic (central) or postrolandic (postcentral) region, whereas the peak value was observed on the posterior parietal veins in the linear model (Figure 5).

**Figure 5.**
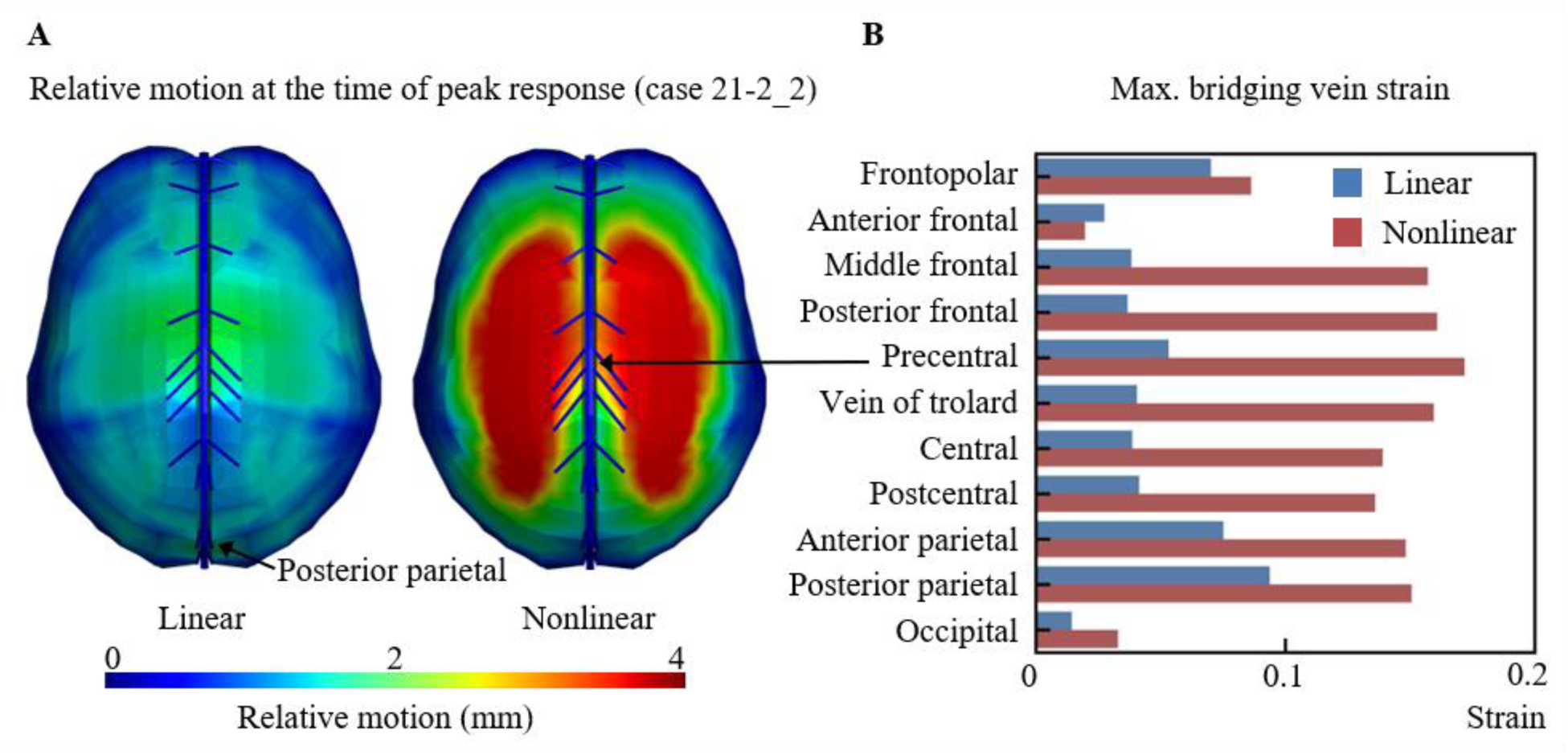
(A) Top view of cortical relative motion distribution at the time of peak response for case 21-2_2; (B) Maximum strain in each pair of bridging veins.

To evaluate the influence of the pial material on brain deformation responses, the element-wise time-accumulated peak of the first principal Green-Lagrange strain (MPS) was analyzed for the cortical region and four deep brain structures (corpus callosum, thalamus, midbrain, and brainstem). For the brain cortex, changing the pia mater from the linear to the nonlinear material relocated the site of peak cortical MPS from the parietal lobe to the frontal lobe in case 21-2_2 (Figure 6A). For the cortical region (Figure 6B), the nonlinear model predicted lower peak MPS than the linear model: 0.22 (8.63% lower) in case 21-2_2, 0.14 (15.1% lower) in case 25-2_1, and 0.16 (11.6% lower) in case 25-2_2. The mean MPS value in the cortical region was reduced by 7.82%, 8.28%, and 5.66% for cases 21-2_2, 25-2_1, and 25-2_2, respectively, when the pia mater was changed from the linear to the nonlinear material.

**Figure 6.**
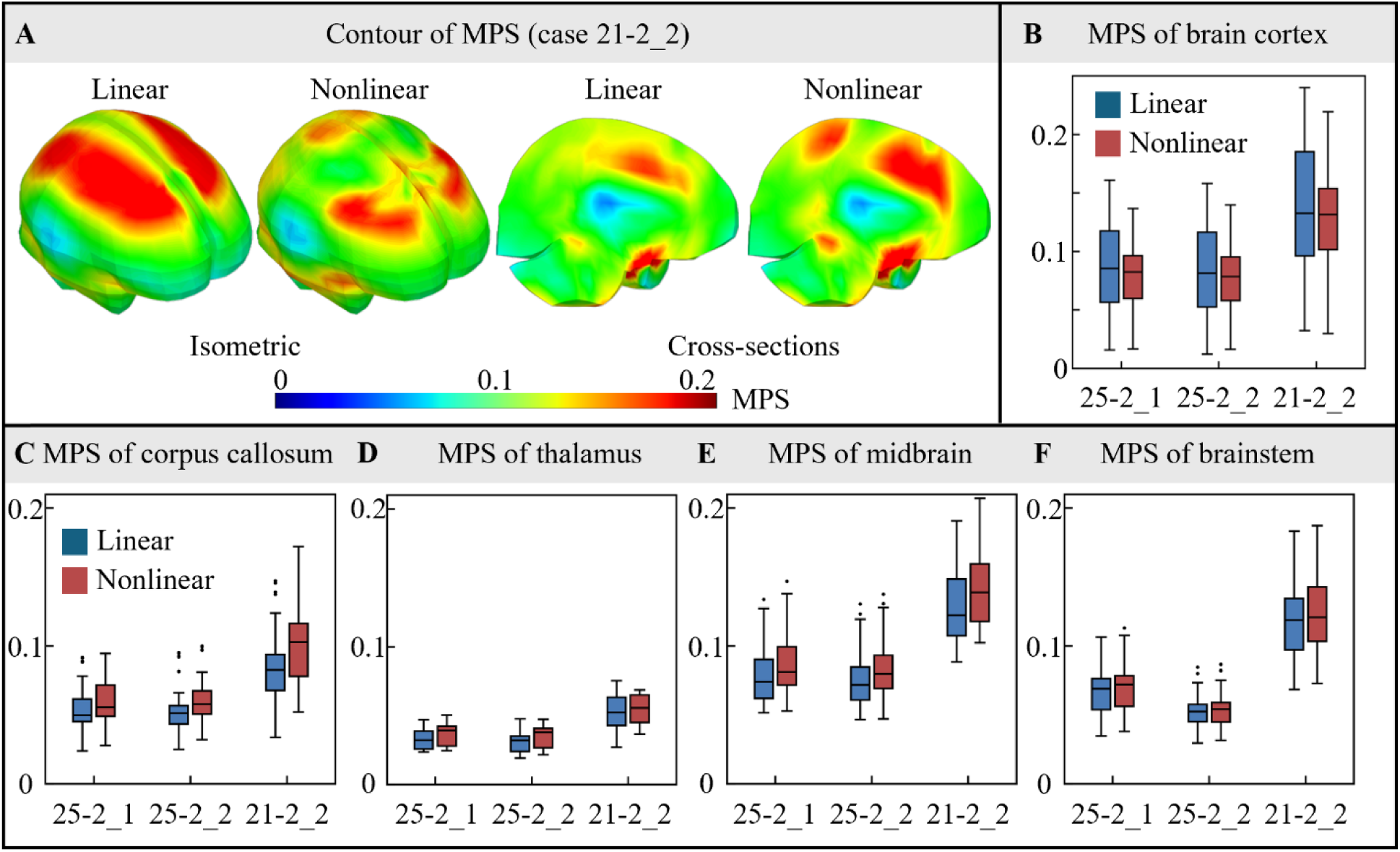
Influence of the pial modeling (linear vs. nonlinear) on the maximum first principal Green-Lagrange strain (MPS) in the brain in three simulated impacts (i.e., cases 25-2_1, 25-2_2, and 21-2_2). (A) Isometric view and sagittal cross-section of MPS distribution of case 21-2_2; (B-F) Comparisons of MPS of the brain elements of different regions: (B) Brain cortex; (C) Corpus callosum; (D) Thalamus; (E) Midbrain; (F) Brainstem. On each box, the central line is the median value, and the upper and lower edges of the box are the 25th and 75^th^ percentile values; the upper and lower edges of the line are the minimum and maximum values, while the outliers are shown as ‘‘·’’ symbol outside the box.

In contrast to the pronounced variations observed in the cortical region, the four deep brain structures exhibited higher MPS values in the nonlinear model than in the linear model (Figure 6C-F). For the peak MPS in case 21-2_2, the nonlinear model increased the values by 17.0%, 8.7%, and 2.2% in the corpus callosum, midbrain, and brainstem, respectively, whereas the thalamic peak decreased from 0.075 to 0.069. The peak MPS increase also held for the two non-rupture cases, ranging from 2.4% (brainstem) to 9.8% (midbrain), with the exception of the thalamus in case 25-2_2 (-0.8%). For the mean MPS in case 21-2_2, the nonlinear model exceeded the linear model by 23.7%, 5.6%, 10.2%, and 3.5% in the corpus callosum, thalamus, midbrain, and brainstem, respectively. This increase held for the two non-rupture cases, with the mean MPS rising by 3.0% (brainstem) to 11.6% (corpus callosum).

## 4. Discussion

The current study evaluated the pia mater’s influence on the prediction of ASDH by applying three experimentally measured loadings with known outcomes of BV conditions to two FE head models with linear or with nonlinear, experimentally based pial material. The results showed that the choice of pia mater modeling strategy altered the magnitude and distribution of the cortical relative motion and brain strain. When interpreting the model-estimated BV strain with the reported BV-SSS ultimate strain, the predicted ASDH occurrence was consistent with the experimental observation only when the pia mater was modeled as a nonlinear material with experimentally based properties. These results verified the hypothesis that the pia mater material modeling affects the prediction outcome of ASDH.

The current finding that changing the pia material from the stiffer linear material to a nonlinear material led to greater cortical motion could be explained by the numerical mechanics of brain-skull contact. This increased compliance allowed interfacial nodes to undergo greater relative tangential motion along the contact surface, resulting in larger cortical relative motion. In the linear model, the tangential displacement of cortical brain nodes was constrained, with a larger proportion of force transmitted to the brain via the brain-skull interface, and hence elevating cortical MPS (Figure 6A and B). In contrast, the nonlinear model permitted greater cortical sliding and lower cortical MPS, redistributing strain away from the surface and transferring the deformation to the deep brain structures. To verify this, we additionally plotted the shear stress of brain cortex, which was higher in the linear model than in the nonlinear model (Figure 7), indicating a large proportion of forces transmitted to the brain via the brain-skull interface. These observations are consistent with the prior studies reporting that enabling cortical sliding at the brain-skull interface extended stress and strain from the surface to internal areas of the brain in FE head models (Kleiven et al. 2002, Zhou et al. 2019b, 2020, Yang et al. 2022).

**Figure 7.**
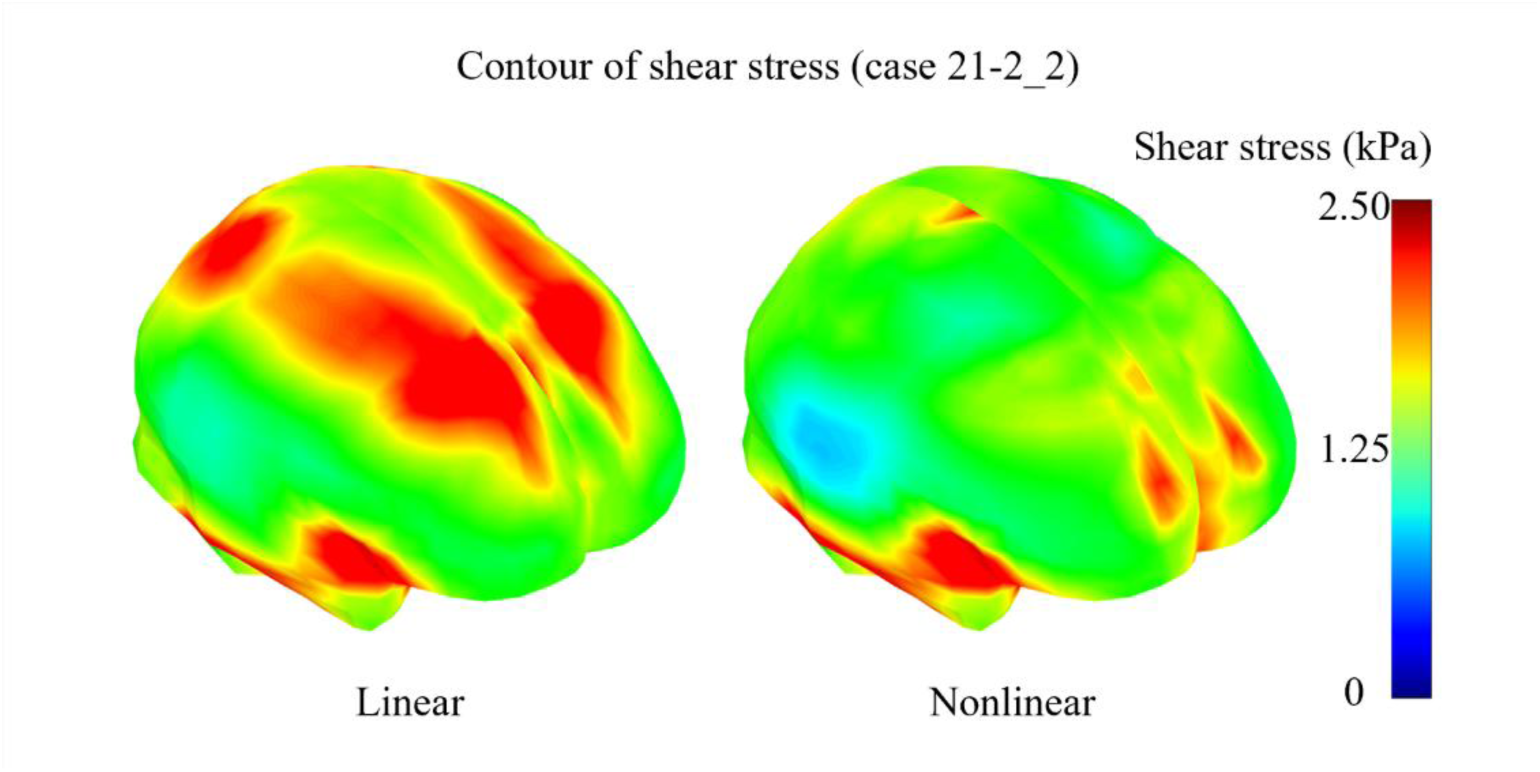
Isometric view of shear stress distribution of case 21-2_2. The nonlinear model (right) produces lower shear stress (e.g., a maximum value of 3.5 kPa and a mean of 1.5 kPa) on the cortical surface than the linear model (left, a maximum value of 3.7 kPa and a mean of 1.8 kPa).

The pia mater itself has received less attention than other aspects of the brain-skull interface, whereas contact algorithms (Kleiven et al. 2002, Joldes et al. 2008, Chafi et al. 2009, Wang et al. 2018, Zhou 2019, Zhou et al. 2019a, b, Yang et al. 2022) and CSF modeling (Chafi et al. 2009, Iwamoto et al. 2015, Zhou et al. 2016) have been studied extensively. The nonlinear mechanical behavior of the pia mater has been treated as part of the meninges rather than represented in isolation (Takhounts et al. 2008). For example, Gu et al. (2012) investigated another FE head model, which excluded the pia mater together with other layers, to demonstrate that meninges act as protective layers for the brain tissue subjected to blast loadings. Beyond impact biomechanics, Wittek et al. (2008) showed that representing the pia mater as a discrete nonlinear membrane was necessary to reproduce stress–strain relationship from experimental data during needle insertion into the brain, yet the influence of nonlinear modeling has not been isolated investigated in ASDH prediction. The present study filled this knowledge gap by isolating the influence of the pial material model (i.e., linear vs. experimentally based nonlinear) and thereby clarified the unique mechanical role of the pia mater in FE head models.

This study suggests that a multivariate approach combining cortical motion and BV strain provides a more mechanistically complete framework for ASDH prediction than a single metric. Compared to the large changes in cortical motion caused by pia mater modeling, BV strain predictions changed only by a small magnitude (Figure 4). Existing FE head models have represented bridging veins using simplified one-dimensional elements [e.g., beam or spring elements (Viano et al. 2005, Kleiven 2007, Cui et al. 2017)], which capture deformation only at discrete nodal locations on the brain surface assuming a straight, spring-like vessel. When large cortical motion occurs away from the anchor nodes, the discrete representation under-registers injury-relevant deformation. For the case known to induce ASDH (i.e., case 21-2_2) simulated by the nonlinear model in this study, the cortical anchor nodes of the BV elements did not coincide with the region of maximum cortical relative motion. Although the model-estimated BV strain fell within the range of the BV-SSS ultimate strain (0.29 ± 0.13) (Monea et al. 2014), and thus indicated possible ASDH, it remained below the range of the BV ultimate strain (0.50 ± 0.16) (Lee et al. 1989). Kapeliotis et al. (2019) compared four models with different BV entry angles and found similar predictive capability, suggesting that BV strain from assuming a straight, spring-like vessel alone may be insufficient to distinguish injury-critical scenarios. A multivariate framework that integrates cortical motion and strain from anatomically detailed models of bridging veins could be better positioned to reflect the complexity of ASDH initiation.

In addition to BV rupture secondary to cortical relative motion, cortical contusion and laceration of the cerebral vessels have also been proposed as candidate instigators of ASDH (Gennarelli et al. 1982). The mechanism of subdural hematoma from sources other than bridging veins should be explored (Mallory 2014). This motivated us to further examine the cortical responses that could be related to ruptured cortical vessels on the surface of the brain. In the present study, we introduced the cortical MPS as a complementary predictor, considering that cerebral vessels are coated and tethered by pia-arachnoid tissue (Alcolado et al. 1988, Monson et al. 2005). For the injury case (21-2_2), the peak cortical MPS (0.24 for linear; 0.22 for nonlinear; Figure 6B) approached the lower bound of the reported cortical artery ultimate strain from autopsy (0.27 ± 0.05) (Monson et al. 2005), whereas the peak cortical MPS in the non-injury cases remained below this range. However, it should be clarified that the experiments of Depreitere et al. (2006) were known to cause BV disruption-induced ASDH. To the best of our knowledge, the loading conditions that produce subdural hematoma from sources other than bridging veins remain unknown.

One limitation that should be acknowledged is the absence of direct experimental validation of cortical surface motion under ASDH-relevant loading conditions. Both computational and experimental evidence indicate that ASDH risk and cortical response are strongly geometry-dependent, underscoring the need for specimen-specific experimental data (Zhou et al. 2019a, Mallory et al. 2026). Tesny (2022) used high-frequency ultrasound to directly measure cortical motion in five postmortem human subjects under controlled impact conditions. Building on such methods, future work could develop personalized FE head models that incorporate subject-specific geometry and validate them against direct cortical motion measurements. In addition, the brain tissue was modeled as an isotropic, homogeneous material in the present study; however, constitutive modeling choices of brain tissue, such as white matter anisotropy and the tension–compression switch schemes in fiber-reinforced model, can also affect predicted brain responses (Li et al. 2026) and could be coupled with nonlinear pial modeling in future work.

## 5. Conclusions

The current study evaluated the consequences of pial material modeling on the prediction of ASDH. The results demonstrated that the choice of material representation (a linear elastic material vs. a nonlinear material with experimentally based properties) led to differences in predicted brain cortical motion and BV strain, highlighting the sensitivity of model predictions to pial material modeling choices. The widely used linear elastic pial material was shown to be an over-stiffening formulation that underestimated the brain cortical response. This study advocates the adoption of experimentally derived nonlinear material for the pia mater, which improves the prediction accuracy of ASDH occurrence.

## Appendix A. rain motion validation of KTH head model with linear and nonlinear pia mater

We validated the KTH head model, incorporating both linear and nonlinear pia mater properties, using the experimental brain motion dataset of Alshareef et al. (2021). In that experiment, six human cadaveric head specimens were instrumented with an array of 32 crystals embedded in the head and brain, 24 of which were implanted into the brain tissue to measure brain motion relative to the skull *in situ*. Each specimen was subjected to head rotational half-sine pulses of three anatomical axes (sagittal, coronal, and axial) with a peak angular velocity of 20 or 40 rad/s and a duration of 30 or 60 ms (Table A1).

For each simulation, the recorded experimental kinematic curves were prescribed at the model’s center of gravity, and the simulated relative brain motions were compared with the marker data. Model performance was quantified using the correlation and analysis (CORA) score (Gehre et al. 2009), defined as the equally weighted average of three sub-ratings (between 0 and 1) that assess the phase, size, and progression of the simulated displacement relative to the experiment. For each 3-D signal, a single representative rating was then obtained by weighting the CORA scores (CORA*_x_*_, *y*, *z*_) by the experimental motion along the three axes (*d_x_*, *d_y_*, *d_z_*), as in Eq. A1 (Davis et al. 2016, Wu et al. 2019).

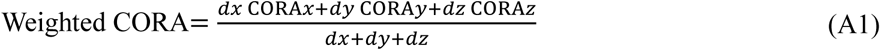

**Table A1.**
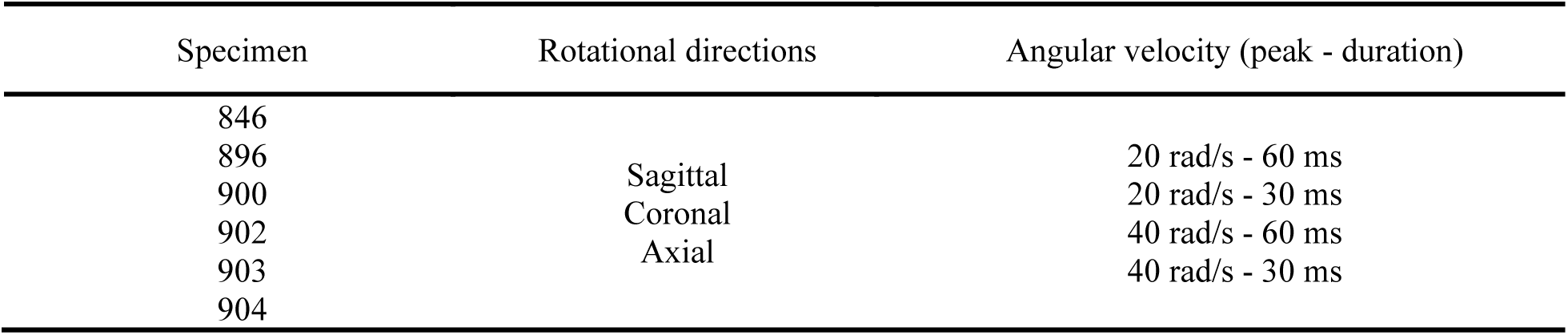
Summary of head kinematics for brain motion validation.

For illustration purposes, validation results of one randomly selected marker in Specimen 846 are shown in Figure A1, of which *d_x_* = 2.25 mm, *d_y_* = 4.92 mm, *d_z_* = 1.06 mm under loading scenario of axial rotation with angular velocity peak of 20 rad/s and a duration of 30 ms. The nonlinear model produced a result closer to the experimental motion than the linear model: the nonlinear vs. linear weighted CORA scores were 0.495 vs. 0.474 for this marker (i.e., 0.570 vs. 0.557 along X, 0.513 vs. 0.488 along Y, and 0.252 vs. 0.233 along Z shown in Figure A1). The same operation was applied to each individual marker, and the marker-specific weighted CORA scores were equally averaged for this illustrative loading case.

**Figure A1.**
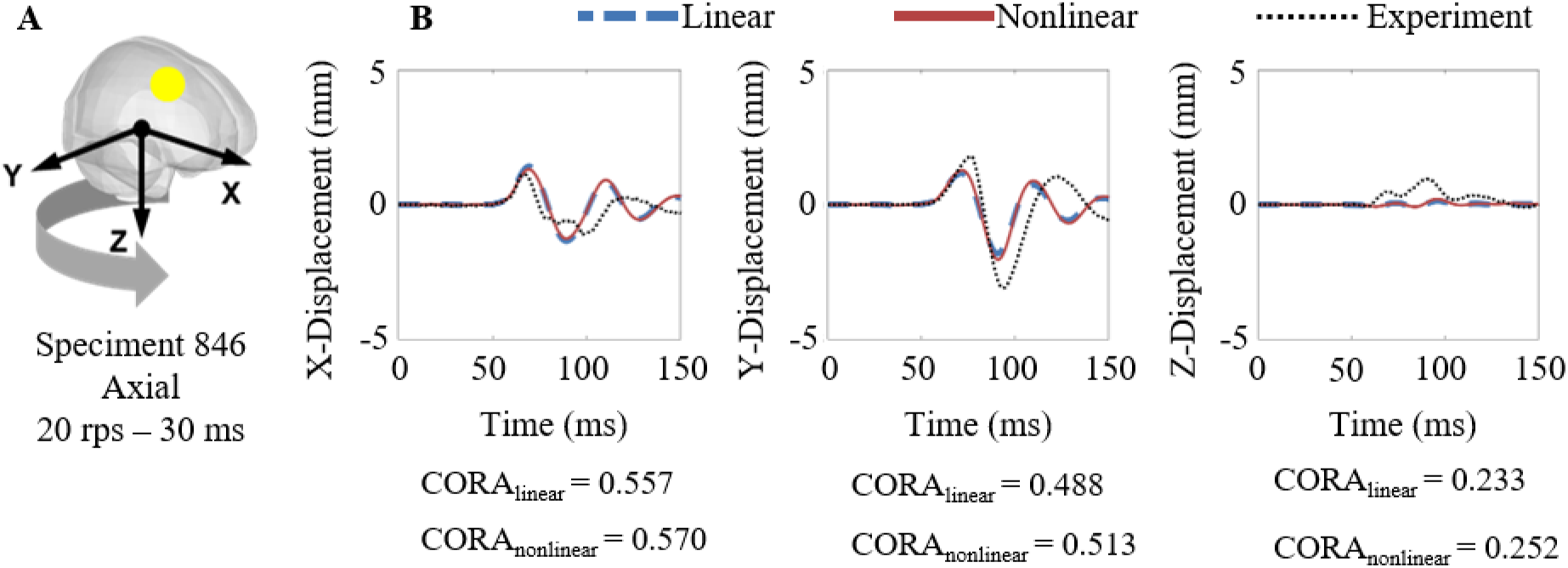
Illustration of one brain motion experiment with one representative marker (in yellow) (A), of which motion results in three anatomical directions (X, Y, Z) are shown in subfigure B with the CORA scores listed.

Across all 72 validation simulations (6 specimens × 12 loading cases), the linear and nonlinear models demonstrated mean weighted CORA scores of 0.522 and 0.528, respectively. Both fall within the range of fair biofidelity (0.44-0.65), according to ISO/TR 9790.

**Figure A2.**
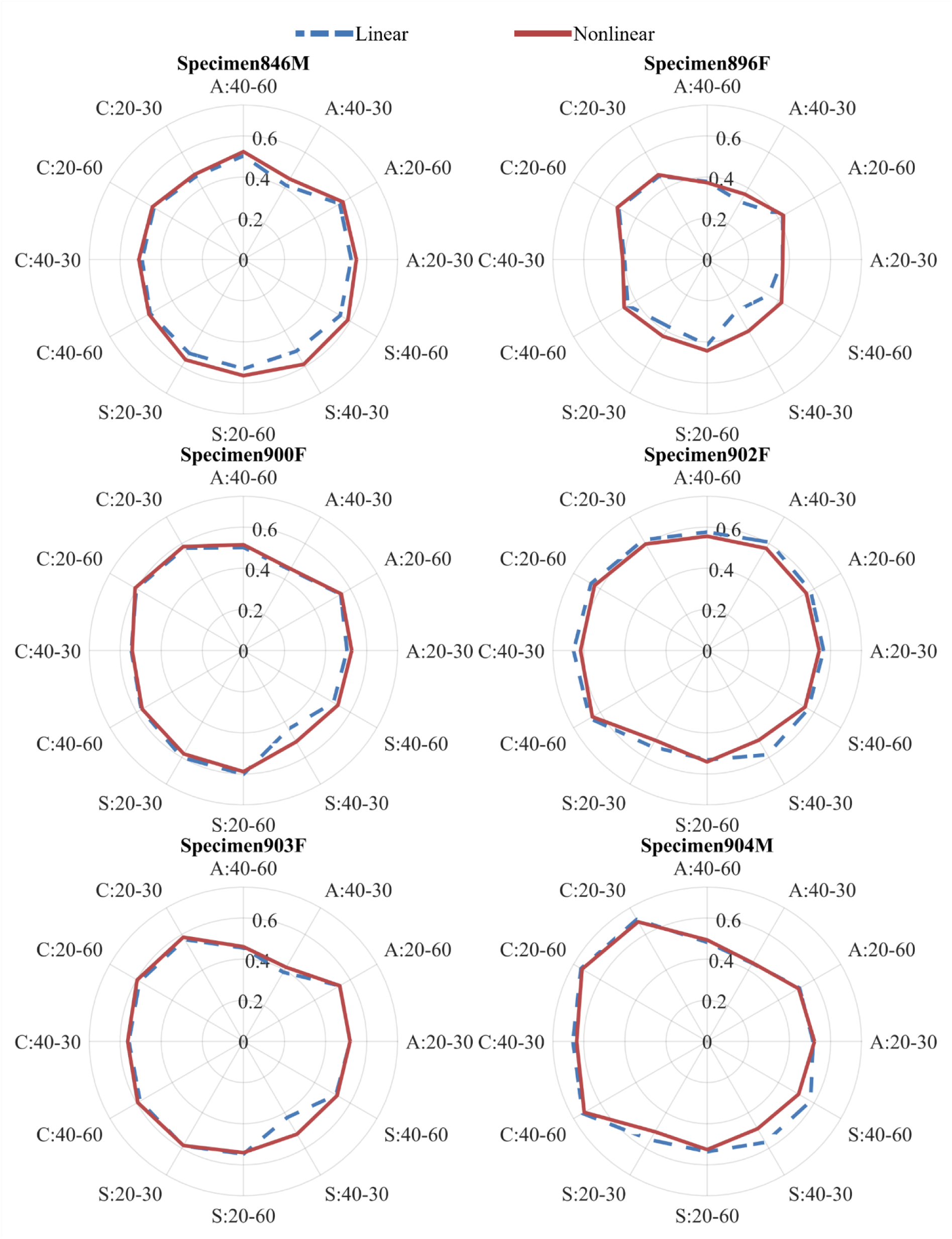
CORA scores of brain motion validation in specimens 846 (top left), 896 (top right), 900 (middle left) 902 (middle right), 903 (bottom left) and 904 (bottom right). In the label of the net plot, the first letter represents the rotational direction (A: Axial, S: Sagittal, C: Coronal), the second number represents the angular velocity peak in the unit of rad/s, and the third number represents the rotational duration in the unit of ms.

## Acknowledgments

This research has received funding from KTH Royal Institute of Technology (Stockholm, Sweden), Swedish Research Council (VR-2024-05848, and VR-2024-02782), and FFI (Strategic Vehicle Research and Innovation), project numbers 2023-00753 and 2024-03635, funded by Vinnova, the Swedish Transport Administration, the Swedish Energy Agency, and the industrial partners. We acknowledge Professor Matthew Panzer from the University of Virginia for providing the experimental data that was used in the current study. The content of this article is solely the responsibility of the authors and does not necessarily represent the official views of the funding agencies. The computational simulations were enabled by resources provided by the National Academic Infrastructure for Supercomputing in Sweden (NAISS) at the center for High Performance Computing (PDC) partially funded by the Swedish Research Council through grant agreement no. 2026/3-281 and 2026/4-723. The authors declare that they have no known competing financial interests or personal relationships that could have appeared to influence the work reported in this paper.

